# Whole-genome analysis of introgression between the spotted owl and barred owl (*Strix occidentalis* and *Strix varia*, respectively; Aves: Strigidae) in western North America

**DOI:** 10.1101/343202

**Authors:** Zachary R. Hanna, John P. Dumbacher, Rauri C.K. Bowie, James B. Henderson, Jeffrey D. Wall

## Abstract

As the barred owl (*Strix varia*; Aves: Strigiformes: Strigidae) expands throughout western North America, hybridization between barred and spotted owls (*Strix varia* and *S. occidentalis*, respectively), if abundant, may lead to genetic swamping of the endangered spotted owl. We analyzed low-coverage, whole-genome sequence data from fifty-one barred and spotted owls to investigate recent introgression between these two species. Although we obtained genomic confirmation that these species can and do hybridize and backcross, we found no evidence of widespread introgression. Plumage characteristics of western *S. varia* that suggested admixture with *S. occidentalis* appear unrelated to *S. occidentalis* ancestry and may instead reflect local selection.

## Introduction

Over the past century, humans have introduced several non-native vertebrate species in western North America into the native range of closely related species and generated moving hybrid swarms. For example, in California, genes of the non-native barred tiger salamander (*Ambystoma tigrinum*) are spreading into the range of the California tiger salamander (*A. californiense*) (Fitzpatrick *et al.* 2009, 2010). In the Flathead River system of Montana and British Columbia, the non-native rainbow trout (*Oncorhynchus mykiss*) is rapidly hybridizing with the native westslope cutthroat trout (*O. clarkii lewisi*) (Muhlfeld *et al.* 2014). In addition to hybridization resulting from intentional introductions of non-native species, changing global climatic conditions and the documented movement of species ranges have led many species to invade novel geographic regions (Parmesan *et al.* 1999; Parmesan 2006) and establish broad contact with related taxa (Rieseberg *et al.* 2007).

The spotted owl (*Strix occidentalis*) is a large wood owl inhabitant of western North American forests. The U.S. Fish and Wildlife Service listed the northern spotted owl (*S. o. caurina*) as “threatened” under the Endangered Species Act (ESA) in 1990 (Thomas *et al.* 1990) and the species remains protected due to continuing population declines (Dugger *et al.* 2015; Davis *et al.* 2016). While researchers considered habitat loss the primary threat to the northern spotted owl in 1990 (Forsman *et al.* 1984; Anderson and Burnham 1992), recent research has confirmed a second major threat to its persistence: the invasion of the congeneric barred owl (*S. varia*) into western North American forests (Dugger *et al.* 2015; Diller *et al.* 2016). Previously inhabiting areas east of the Rocky Mountains and Great Plains (Mazur and James 2000), the barred owl has expanded its range to western North America over the last 50-100 years (Dark *et al.* 1998; Livezey 2009a, 2009b). At present, sympatric populations of spotted and barred owls exist from British Columbia to southern California (Taylor and Forsman 1976; Haig *et al.* 2004; Livezey 2009a).

Spotted and barred owls are approximately 13.9% divergent in the mitochondrial control region (Haig *et al.* 2004), 10.74% divergent in non-tRNA mitochondrial genes (Hanna *et al.* 2017a), and 0.7% divergent across the nuclear genome (Hanna *et al.* 2017c). Barred and spotted owls hybridize and backcross (Haig *et al.* 2004; Kelly and Forsman 2004; Funk *et al.* 2007), with heterospecific matings and F_1_ hybrids commonly reported in areas where barred owls are rare and spotted owls common (Kelly and Forsman 2004). Western barred owl specimens in museum collections display striking morphological variation. Birds from the eastern Klamath Mountains in Siskiyou County, California, have darker plumage overall, more spotting on the belly, and are smaller than barred owls from the Coast Range (Figure 1 and Figure S1). These differences suggest either local selection for this phenotype or possible introgression of spotted owl genes. Hybridization of these species creates a potential for a loss of biodiversity in western North America due either to replacement of the spotted owl by the barred owl or to collapse of the boundaries of the two species (Huxel 1999).

**Figure 1.**
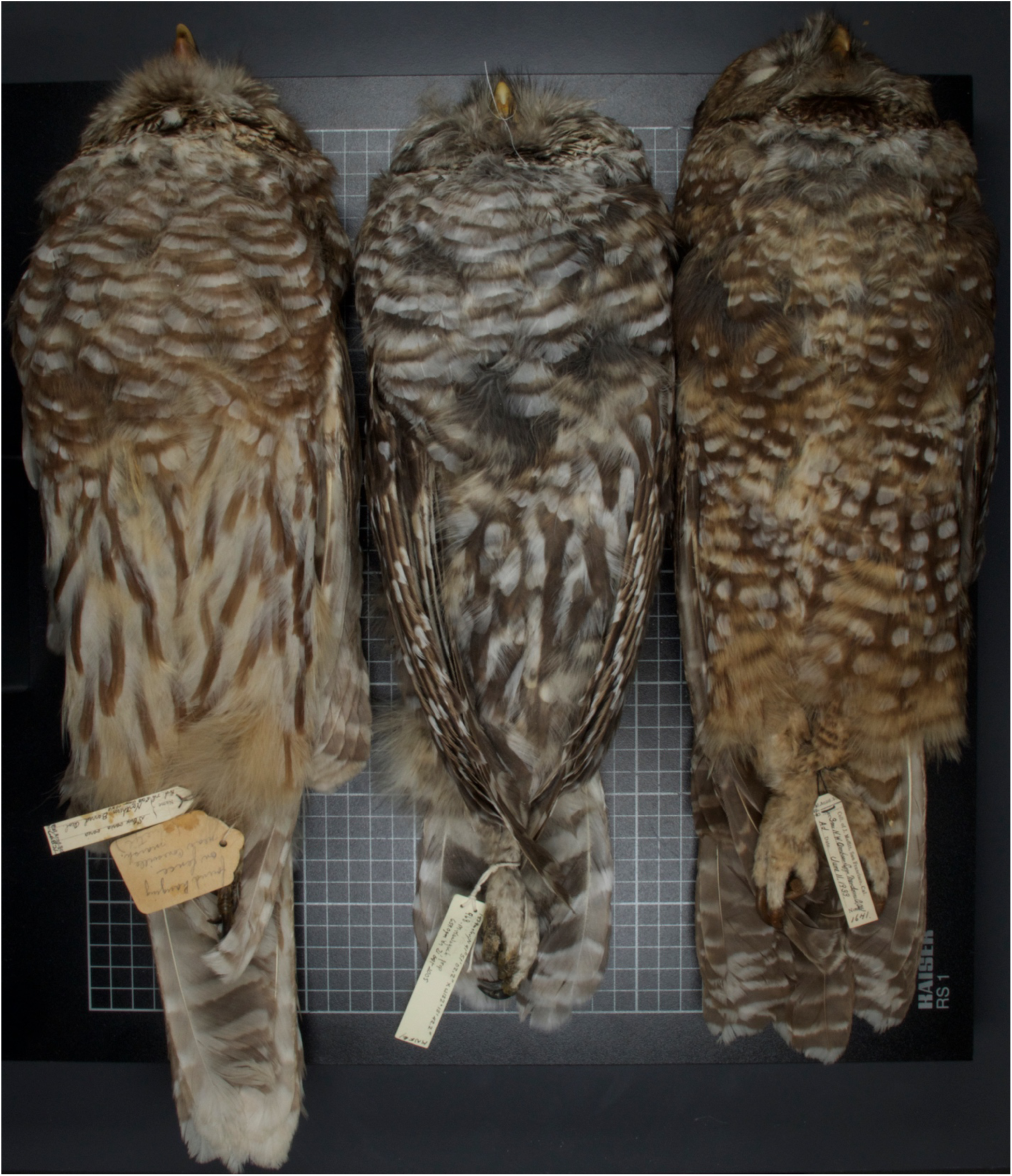
Comparison of eastern barred owl, Siskiyou County barred owl, and northern spotted owl plumages. This image displays the darker ventral plumage of a *Strix varia* collected in Siskiyou County, California compared with that of typical *S. varia* and *S. occidentalis caurina* individuals. On the left is the ventral plumage of a *Strix varia* from eastern North America. In the center is a *S. varia* from Siskiyou County, California. On the right is a *S. occidentalis caurina* from northern California. Author Z.R.H. took this photograph.

For this study, we obtained fifty-one low-coverage whole-genome sequences (median 0.723X coverage) from barred and spotted owls sampled outside and across their contact zone in western North America (Figure 2). We utilized available medium and high-coverage whole-genome sequences from an eastern barred owl (15.549X coverage) and a pre-contact spotted owl (60.815X coverage) to identify variant sites fixed between the barred and spotted owl. For each low-coverage individual, we determined the genome-wide average ancestry and searched for windows of ancestry that were outliers from the average to detect rare, introgressed regions. We used these data to identify the extent of introgression between the barred and spotted owl in western North America.

**Figure 2.**
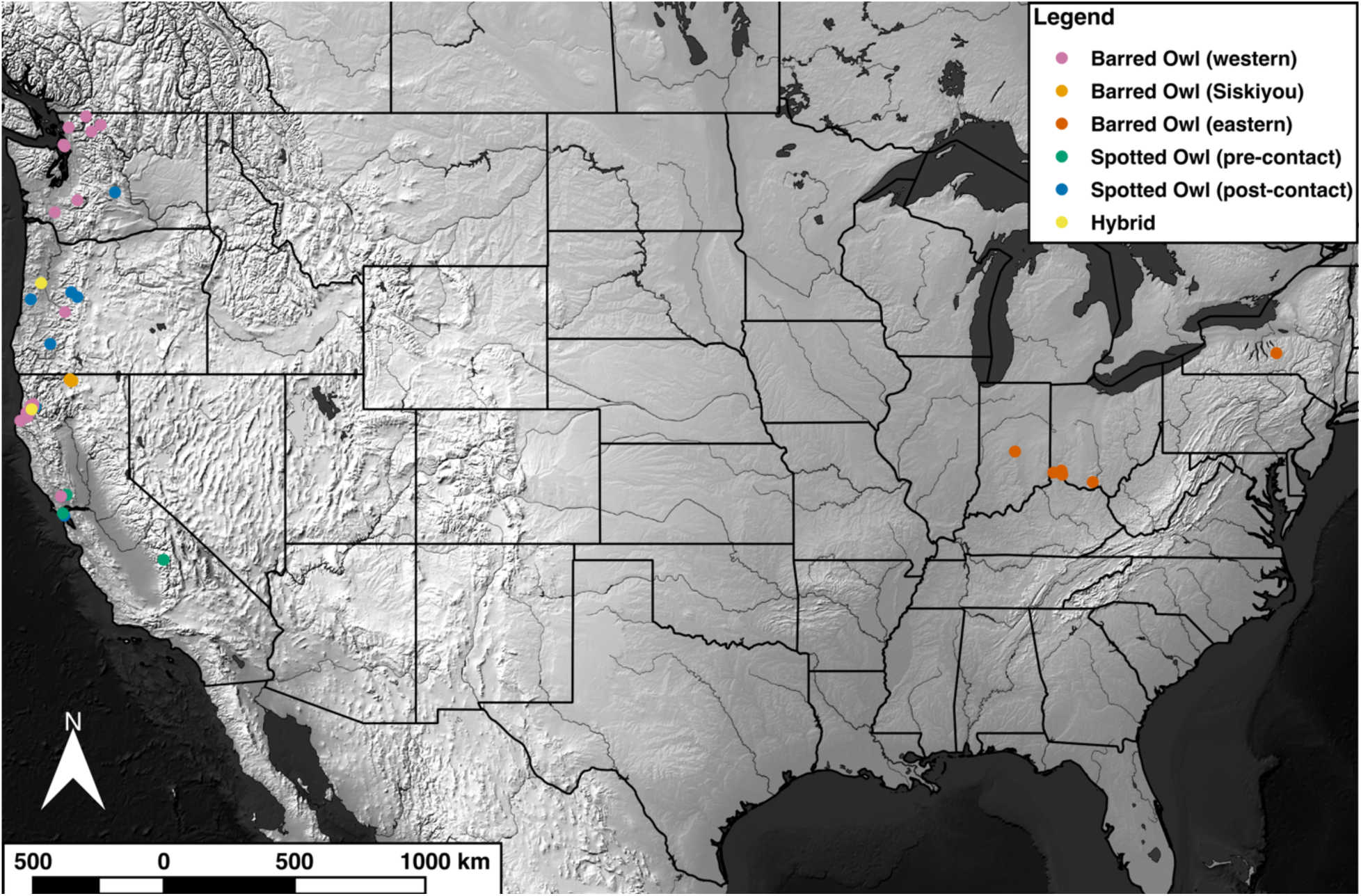
Sample map. This map displays the sampling locations of all of the *Strix* specimens included in this study.

## Methods

### Samples

We obtained fifty-one samples from museum collections that included eleven *Strix occidentalis* samples (two samples predated contact with *S. varia*), thirty-eight *S. varia* samples (including five from eastern North America), and two samples identified by other researchers as probable hybrid *S. varia* × *occidentalis* individuals (Tables S1-S2). We mapped the samples using QGIS version 2.18.2 (Quantum GIS Development Team 2017) with raster and vector files from Natural Earth (http://www.naturalearthdata.com; accessed 2017 Oct 1) (File S1 section 1.1).

### Sequence data

Hybridization of *Strix varia* and *S. occidentalis* has previously been investigated using a set of four microsatellite (Funk *et al.* 2007) and fourteen amplified fragment-length polymorphism (Haig *et al.* 2004) markers, which the authors found useful for diagnosing F_1_ and F_2_ hybrids (Haig *et al.* 2004; Funk *et al.* 2007). We utilized a whole-genome sequencing approach to enable us to detect any introgression that has taken place over the last 50-70 years that *S. varia* and *S. occidentalis* have been in contact in western North America (Taylor and Forsman 1976; Livezey 2009a).

We utilized whole genome sequencing data from a previous study (Hanna *et al.* 2017c) for our reference pre-contact *Strix occidentalis* and eastern *S. varia* samples (NCBI Sequence Read Archive (SRA) run accessions SRR4011595, SRR4011596, SRR4011597, SRR4011614, SRR4011615, SRR4011616, SRR4011617, SRR4011618, SRR4011619, and SRR4011620 for *S. occidentalis* sample CAS:ORN:98821; SRR5428115, SRR5428116, and SRR5428117 for *S. varia* sample CNHM<USA-OH>:ORNITH:B41533, hereafter referred to as CNHMB41533). We prepared whole genome libraries for fifty-one additional (i.e. non-reference) *Strix* samples using a Nextera DNA Sample Preparation Kit (Illumina) and obtained paired-end sequences from a HiSeq 2500 (Illumina) (File S1 section 1.2) resulting in coverage ranging from 0.02-6.41X after filtering.

### Alignment and filtering

For the sequence data of the reference samples *Strix occidentalis* CAS:ORN:98821 and *S. varia* CNHMB41533, which Hanna et al. (2017c) generated for their study, we followed the processing methods described in Hanna et al. (2017c) and used the data here in its final, processed form. For all other samples we used Trimmomatic version 0.32 (Bolger *et al.* 2014) to remove adapter sequences and perform quality trimming of all of the low-coverage, short-read data (File S1 section 1.3). We used BWA-MEM version 0.7.12-r1044 (Li 2013) to align the processed reference and low-coverage sequences to the repeat-masked *S. o. caurina* genome “StrOccCau_1.0_nuc_masked” (Hanna *et al.* 2017d, 2017c). We merged the alignments, sorted the alignments, and marked duplicate sequences using Picard version 1.104 (http://broadinstitute.github.io/picard; accessed 2017 Oct 1) (File S1 section 1.4.1-1.4.2). We filtered the alignment files to only retain alignments of high quality using the Genome Analysis Toolkit (GATK) version 3.4-46 PrintReads tool (McKenna *et al.* 2010; DePristo *et al.* 2011; Van der Auwera *et al.* 2013; GATK Dev Team 2017) (File S1 section 1.4.3).

### Variant calling and filtering

We called variants using the GATK version 3.4-46 UnifiedGenotyper tool (McKenna *et al.* 2010; DePristo *et al.* 2011; Van der Auwera *et al.* 2013) with the alignment files for all samples included as simultaneous inputs (File S1 section 1.5.1). We used the vcf_qual_filter.sh script from SPOW-BDOW-introgression-scripts version 1.1.1 (Hanna *et al.* 2017b) to exclude indels and low genotyping quality sites while retaining only biallelic sites where CAS:ORN:98821 (the source of the StrOccCau_1.0_nuc_masked reference genome) was homozygous for the reference allele and CNHMB41533, the *Strix varia* reference sample, was homozygous for the alternative allele (File S1 section 1.6.1). Of the remaining variable sites, we excluded those with excessively high coverage [greater than the mean plus five times the standard deviation (σ), as recommended by the GATK documentation (https://software.broadinstitute.org/gatk/guide/article?id=3225; accessed 2017 Oct 1)] (File S1 section 1.6.2). We used DP_means_std_dev.sh from SPOW-BDOW-introgression-scripts version 1.1.1 to calculate the mean and standard deviation (σ) of the depth of coverage for each sample across the final set of variant sites.

### Ancestry and diversity analyses

For each sample at each of the final variant sites, we calculated a percentage spotted owl ancestry, which was the percentage of the coverage that supported the CAS:ORN:98821 (the *Strix occidentalis* reference sequence) allele. We calculated the mean and standard deviation of the spotted owl ancestry of each sample across all variant sites (File S1 section 1.6.3). We tested for significant differences between the mean spotted owl ancestries in populations using Welch’s *t*-test (Welch 1947) as the populations had unequal numbers of samples and then applied a Bonferroni adjustment (Dunn 1961) when we evaluated significance (File S1 section 1.6.4).

We estimated the probabilities of observing an introgressed region greater than 50,000 nt, 100,000 nt, or 150,000 nt in length if *Strix varia* and *S. occidentalis* hybridized in 1945, approximately the earliest date of their potential contact (Livezey 2009a), using the formula from Racimo et al. (2015). For the recombination rate, we used 1.5 centimorgans/million nucleotides (cM/Mnt), which Backström et al. (2010) estimated for the zebra finch (*Taeniopygia guttata*). For the number of generations since the earliest potential date of hybridization, we assumed a generation time of two years (Gutiérrez *et al.* 1995; Mazur and James 2000) even though *S. o. caurina* is able to breed in its first year and others have used ten years as the generation time for *S. o. caurina* (Noon and Biles 1990; USDA Forest Service 1992). With that generation time, approximately thirty-five generations have potentially elapsed since the two species first contacted in 1945 and 2014, the date of our most recent sample.

In order to probe for further for evidence of introgression in the samples that did not appear as hybrids from their genome-wide average spotted owl ancestry, we attempted to identify regions that were outliers from the genome-wide ancestry average by conducting a sliding window analysis. We examined adjacent windows of 50,000 nucleotides (nt) where a sample had data for at least ten variant sites within that window and calculated the average spotted owl ancestry for the window. We assumed that, if a region was introgressed from the other species, the average should be close to 0.5. Thus, in samples with an average genome-wide ancestry close to 0, we called a window an outlier if the average spotted owl ancestry was >= 0.4. Inversely, in samples with an average genome-wide ancestry close to 1, we called a window an outlier if the average spotted owl ancestry was <= 0.6 (File S1 sections 1.6.5-1.6.6).

In order to estimate the genome-wide diversity harbored by *Strix varia* and *S. occidentalis* populations, we considered all biallelic variant sites (not just those fixed between our *S. varia* and *S. occidentalis* references) and calculated π_Within_, the number of nucleotide differences within populations, and π_Between_, the number of nucleotide differences between populations using the countFstPi script from SPOW-BADO-introgression-scripts (Hanna *et al.* 2017b). We also used countFstPi to calculate the fixation index (*F_ST_*) (Hudson *et al.* 1992) in order to estimate the differentiation of *S. varia* and *S. occidentalis* populations (File S1 section 1.6.7).

### Data availability

Raw whole genome sequences are available from the NCBI Sequence Read Archive (SRA) run accessions SRR4011595-SRR4011597, SRR4011614-SRR4011620, SRR5428115- SRR5428117, SRR6026668, SRR6032894-SRR6032902, SRR6032904-SRR6032907, and SRR6032910-SRR6033014. See Table 1 for the specific accessions corresponding with each sample.

**Table 1.** Genomic sequence data details for each sample.

| Voucher Specimen Identifier | Other Sample Identifier | Sample Set | SRA ACCN |
| --- | --- | --- | --- |
| CAS:ORN:98821 | Sequoia | N/A | SRR4011595, SRR4011596, SRR4011597, SRR4011614, SRR4011615, SRR4011616, SRR4011617, SRR4011618, SRR4011619, SRR4011620, |
| CNHM<USA-OH>:ORNITH:B41533 | CMCB41533 | N/A | SRR5428115, SRR5428116, SRR5428117 |
| CAS:ORN:87569 | CAS87569 | 1 | SRR6032959 |
| CAS:ORN:92982 | ASG007 | 1 | SRR6032957 |
| CAS:ORN:95475 | MK994 | 1 | SRR6032939 |
| CAS:ORN:95789 | JMR920 | 1 | SRR6032933 |
| CAS:ORN:95790 | ASG037 | 1 | SRR6032960 |
| CAS:ORN:95964 | MEF457 | 1 | SRR6026668 |
| CAS:ORN:97181 | MK1020 | 1 | SRR6032934 |
| CNHM<USA-OH>:ORNITH:B40819 | CMCB40819 | 1 | SRR6032951 |
| CNHM<USA-OH>:ORNITH:B40824 | CMC40824 | 1 | SRR6032952 |
| CNHM<USA-OH>:ORNITH:B41566 | CMCB41566 | 1 | SRR6032935 |
| CUMV:Bird:51478 | CU51478 | 1 | SRR6032936 |
| MVZ:Bird:189508 | ZRH455 | 1 | SRR6032920 |
| UWBM:Bird:62061 | UWBM62061 | 1 | SRR6032940 |
| UWBM:Bird:76815 | UWBM76815 | 1 | SRR6032937 |
| UWBM:Bird:91379 | UWBM91379 | 1 | SRR6032938 |
| UWBM:Bird:91382 | UWBM91382 | 1 | SRR6032931 |
| UWBM:Bird:91408 | UWBM91408 | 1 | SRR6032932 |
| CAS:ORN:92979 | MK968 | 2 | SRR6032898, SRR6032899, SRR6032916 |
| CAS:ORN:92980 | MK987 | 2 | SRR6032914, SRR6032915, SRR6032917 |
| CAS:ORN:92981 | MEF404 | 2 | SRR6032941, SRR6032945, SRR6032946 |
| CAS:ORN:95476 | MK998 | 2 | SRR6032910, SRR6032912, SRR6032913 |
| CAS:ORN:95477 | ASG017 | 2 | SRR6032902, SRR6032904, SRR6032905 |
| CAS:ORN:97049 | LCW491 | 2 | SRR6032943, SRR6032944, SRR6032950 |
| CAS:ORN:97052 | LCW443 | 2 | SRR6032947, SRR6032948, SRR6032949 |
| CAS:ORN:97174 | MEF432 | 2 | SRR6032894, SRR6032895, SRR6032942 |
| CAS:ORN:97175 | MK1012 | 2 | SRR6033011, SRR6033013, SRR6033014 |
| CAS:ORN:97176 | JPD386 | 2 | SRR6032926, SRR6032927, SRR6032928 |
| CAS:ORN:97177 | MEF435 | 2 | SRR6032896, SRR6032897, SRR6033012 |
| CAS:ORN:97201 | LCW405 | 2 | SRR6032925, SRR6032929, SRR6032930 |
| CAS:ORN:97815 | Hoopa20005 | 2 | SRR6032900, SRR6032906, SRR6032907 |
| CAS:ORN:97816 | Hoopa20018 | 2 | SRR6032924, SRR6032961, SRR6032962 |
| CAS:ORN:97818 | Hoopa20011 | 2 | SRR6032901, SRR6032965, SRR6032966 |
| CAS:ORN:97819 | Hoopa20019 | 2 | SRR6032921, SRR6032922, SRR6032923 |
| CAS:ORN:97820 | Hoopa20017 | 2 | SRR6032967, SRR6032968, SRR6032970 |
| CAS:ORN:97822 | Hoopa20014 | 2 | SRR6032963, SRR6032964, SRR6032969 |
| CAS:ORN:98171 | ZRH962 | 2 | SRR6032955, SRR6032956, SRR6032958 |
| CAS:ORN:98198 | ZRH602 | 2 | SRR6032992, SRR6032997, SRR6032998 |
| CAS:ORN:99315 | ZRH604 | 2 | SRR6032995, SRR6032996, SRR6032999 |
| CAS:ORN:99320 | ZRH607 | 2 | SRR6032953, SRR6032954, SRR6033000 |
| CAS:ORN:99423 | NSO138799040 | 2 | SRR6032911, SRR6032918, SRR6032919 |
| CAS:ORN:99425 | NSO168709365 | 2 | SRR6032988, SRR6032989, SRR6032990 |
| UWBM:Bird:53433 | UWBM53433 | 2 | SRR6032985, SRR6032986, SRR6032987 |
| UWBM:Bird:65055 | UWBM65055 | 2 | SRR6032982, SRR6032983, SRR6032984 |
| UWBM:Bird:67015 | UWBM67015 | 2 | SRR6032981, SRR6033003, SRR6033004 |
| UWBM:Bird:74078 | UWBM74078 | 2 | SRR6033005, SRR6033006, SRR6033007 |
| UWBM:Bird:79007 | UWBM79007 | 2 | SRR6033008, SRR6033009, SRR6033010 |
| UWBM:Bird:79049 | UWBM79049 | 2 | SRR6032972, SRR6033001, SRR6033002 |
| UWBM:Bird:79141 | UWBM79141 | 2 | SRR6032971, SRR6032973, SRR6032974 |
| UWBM:Bird:91380 | UWBM91380 | 2 | SRR6032975, SRR6032976, SRR6032978 |
| UWBM:Bird:91392 | UWBM91392 | 2 | SRR6032977, SRR6032979, SRR6032980 |
| UWBM:Bird:91393 | UWBM91393 | 2 | SRR6032991, SRR6032993, SRR6032994 |

The “Specimen Identifier” column provides the voucher specimen codes. The “Other Sample Identifier” column provides an abbreviated sample code. Column “Sample Set” refers to the round of sequencing that produced the sequence data for a given sample. The main and supplemental methodology sections provide details of the production of these two sets of
495 sequence data. Column “SRA ACCN” provides NCBI Sequence Read Archive (SRA) run accessions in which the raw sequences for each sample are archived.

## Results

After filtering, the final set of variable sites fixed between the *Strix varia* and *S. occidentalis* reference individuals included 5,816,692 sites. The median genome coverage per individual was 0.723X (Table S3). Except for the two putative hybrid samples that we included as a test of our methodology, the genome-wide average spotted owl ancestry for all samples was close to either 0 or 1, indicating that they were either pure *S. varia* or *S. occidentalis*, respectively (Figure 3 and Table S3). A genome-wide average spotted owl ancestry of 0.538 confirmed the F_1_ hybrid (*S. varia* × *occidentalis*) identity of a sample from Humboldt County, California. We calculated a spotted owl ancestry of 0.359 for the second hybrid sample from Benton County, Oregon, which suggested that this individual was likely a F_2_ hybrid (F_1_ × *S. varia* backcross).

**Figure 3.**
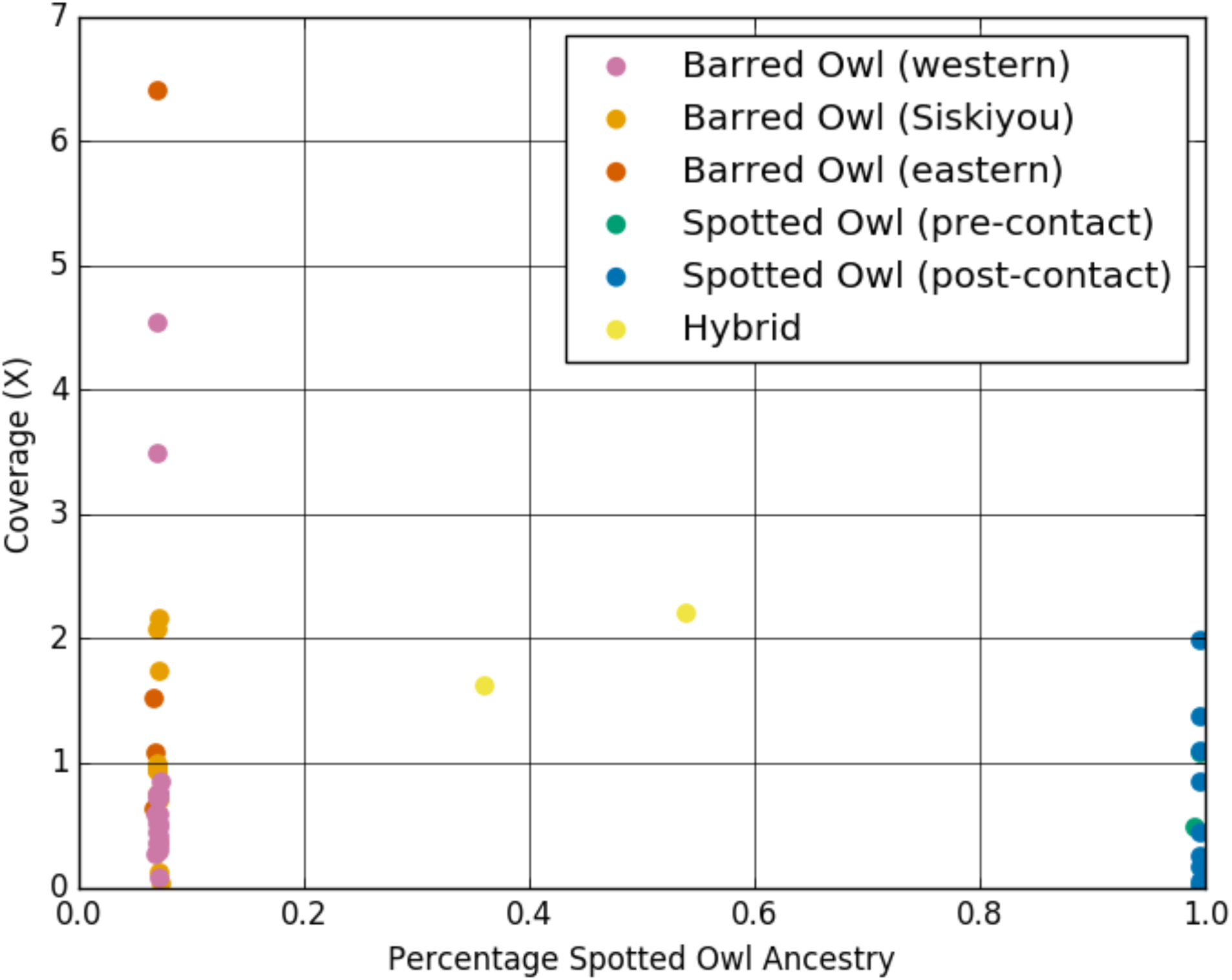
Plot of coverage versus genome-wide average spotted owl ancestry. The average spotted owl (*Strix occidentalis*) ancestry of all of the samples for which we collected low-coverage, whole-genome sequence data. We plotted DNA sequence coverage on the y-axis to display that the average percentage of spotted owl ancestry was independent of the amount of coverage for a given sample.

The mean genome-wide spotted owl ancestry of the Siskiyou County *Strix varia* population was 0.0696 whereas the mean was 0.0699 for the rest of the western *S. varia* (Table S4). There was no significant difference in spotted owl ancestry between these two populations (Table S5). When we combined all *S. varia* from western North America together (0.0698 mean spotted owl ancestry) and compared their spotted owl ancestry with that of the eastern *S. varia* (0.0676 mean spotted owl ancestry), we found no significant difference in ancestry between the western and eastern *S. varia* after applying a Bonferroni adjustment (Tables S4 and S5). There was also no significant difference in spotted owl ancestry between *S. occidentalis* individuals sampled from populations not in contact with *S. varia* and those from populations already in contact with *S. varia* (mean ancestries of 0.9930 and 0.9952, respectively) (Tables S4 and S5).

The average spotted owl ancestry in the *Strix varia* samples ranged from approximately 6.55-7.28% greater than the 0% value at which our methodology set the reference *S. varia* (Table S3). The *S. occidentalis* samples ranged from approximately 0.43-0.94% less than the 100% value for the reference *S. occidentalis*. The standard deviation in the *S. varia* samples was consistently more than two times greater than the standard deviation in the *S. occidentalis* samples. The genome-wide average spotted owl ancestry values for the *Strix varia* individuals deviated more from those of the reference *S. varia* than did the *S. occidentalis* individuals from the *S. occidentalis* reference due to the greater amount of genetic variation within *S. varia* (Hanna *et al.* 2017c). It was evident that the sites fixed between our reference *S. varia* and *S. occidentalis* samples were not fixed across *S. varia* and *S. occidentalis*. Further high-coverage sequencing of whole-genomes for both species will help to more clearly identify the fixed genetic differences between the two species.

Based upon an estimate of thirty-five generations as the maximum number of generations since contact of *Strix varia* and *S. occidentalis* (Gutiérrez *et al.* 1995; Mazur and James 2000; Livezey 2009a) and the recombination rate of *Taeniopygia guttata* (Backström *et al.* 2010), we estimated that the probability of observing a track > 50,000 nt resulting from hybridization during the initial contact of *S. varia* and *S. occidentalis* was 97.41%, the probability of observing a track > 100,000 nt was 94.89%, and the probability of observing a track > 150,000 nt was 92.43%.

Of the forty-nine samples for which we conducted an outlier window analysis, we detected outlier windows in thirty-nine samples (79.6%). Across all samples, we detected 316 outlier windows of length 50,000 nt, forty-one of length 100,000 nt, and only three of length 150,000 nt and none exceeded this length (Figure S2). In all samples the outlier windows represented < 1.01% of the analyzed windows. For thirty-six of the thirty-nine samples with outliers, the number of outlier windows was < 0.08% of the analyzed windows. There were three samples for which the outlier windows represented between 0.1% and 1.01% of the analyzed windows. However, the increased proportion of outlier windows in these samples appeared to be related to exceptionally low sequence coverage as these three *Strix varia* samples had the lowest coverage (0.036-0.118X) and, consequently, the fewest number of analyzed windows of any of the samples in which we detected outlier windows (Figure S3). A *S. occidentalis* sample with 0.017X coverage was the only sample with lower coverage than those three, but our analyses did not recover any outlier windows for it.

We found little evidence of differentiation between the Siskiyou *Strix varia* and the other western *S. varia*, recovering a low *F_ST_* (0.008) and very similar levels of nucleotide diversity in the two populations (Table 2). Similar levels of nucleotide diversity also exist in the *S. varia* populations from western and eastern North America. We additionally estimated a low *F_ST_* value (0.051) between western and eastern *S. varia*, which suggests a low level of differentiation between these populations. *Strix occidentalis* populations pre and post-contact with *S. varia* exhibited similar levels of nucleotide diversity and appeared weakly differentiated (*F_ST_* = 0.022). We estimated approximately 14X greater nucleotide diversity in *S. varia* than *S. occidentalis* and a high level of divergence (*F_ST_* = 0.833) between the species.

**Table 2.** Nucleotide diversity and fixation index statistics calculated for various population comparisons.

| **Population 1** | **Population 2** | **$\pi_{\text{Within Pop 1}}$** | **$\pi_{\text{Within Pop 2}}$** | **$\pi_{\text{Between}}$** | **$F_{ST}$** |
| --- | --- | --- | --- | --- | --- |
| Western Barred Owls | Siskiyou Barred Owls | 2.097E-03 | 2.068E-03 | 2.100E-03 | 0.008 |
| Western Barred Owls | Eastern Barred Owls | 2.119E-03 | 2.228E-03 | 2.291E-03 | 0.051 |
| Siskiyou Barred Owls | Eastern Barred Owls | 2.066E-03 | 2.203E-03 | 2.259E-03 | 0.055 |
| All Western Barred Owls | Eastern Barred Owls | 2.128E-03 | 2.242E-03 | 2.301E-03 | 0.051 |
| All Barred Owls | All Spotted Owls | 2.202E-03 | 1.572E-04 | 7.052E-03 | 0.833 |
| Spotted Owls (pre-contact) | Spotted Owls (post) | 1.073E-04 | 9.998E-05 | 1.060E-04 | 0.022 |

500 The $\pi_{\text{Within}}$ statistic signifies the average number of pairwise differences between two individuals sampled from the same population. The $\pi_{\text{Between}}$ statistic denotes the average number of pairwise differences between two individuals sampled from different populations (Populations 1 and 2). “Pop 1” and “Pop 2” refer to Population 1 or 2 from columns 1 and 2, respectively. The “All Western Barred Owls” population is a combination of the “Western Barred Owls” and “Siskiyou 505 Barred Owls” populations. The “Spotted Owls (pre-contact)” and “Spotted Owls (post)” populations indicate *Strix occidentalis* from populations not in contact or in contact with *S. varia*, respectively.

## Discussion

Our genome-wide average spotted owl ancestry analysis confirmed that our two positive control hybrids from Humboldt County, California, and Benton County, Oregon, were an F_1_ and F_2_ (F_1_ × *Strix varia*) backcross, respectively. Apart from those hybrids, our genome-wide average spotted owl ancestry analysis indicated that all individuals were either pure *S. occidentalis* or pure *S. varia* (Figure 3 and Table S3). Our global analysis found no evidence for admixture, but by averaging over the whole genome we may have missed rare, introgressed regions. Therefore, we implemented a sliding window approach to determine whether any such regions existed in our data. Scanning for ancestry windows that were outliers from a given individual’s genome-wide average ancestry using a sliding window approach corroborated the genome-wide average results and provided no evidence of introgression between *Strix varia* and *S. occidentalis* within the past 50-70 years of their contact in western North America (Taylor and Forsman 1976; Livezey 2009a). Although our test found short windows of outlier ancestry, these represented a small proportion of the total windows analyzed for each individual. Thus, we can confidently exclude the possibility of introgression within the past ten generations. Hybridization that has occurred in the last thirty-five generations (assuming a generation time of two years for both species and erring conservatively on the side of overestimating the maximum number of generations of contact) should have yielded much longer outlier blocks than we found. Even with this conservative estimate, there is a > 97% probability of introgressed regions being larger than the 50,000 nt windows that we used to check for potential introgression and a > 92% probability of the introgressed regions being larger than the 150,000 nt length of the longest outlier window that we detected with our sliding window analysis.

Since *Strix varia*’s zone of contact with *S. occidentalis* in western North America began in British Columbia and expanded southward to the southern Sierra Nevada, California (Taylor and Forsman 1976; Haig *et al.* 2004; Livezey 2009a), we expected *S. varia* individuals in the southern portion of the zone of sympatry to have the highest chance of being admixed. With this prediction in mind, we focused our sampling on *S. varia* populations in California (Figure 1) and targeted our sampling to include the morphologically anomalous western *S. varia* population in Siskiyou County, California. It is notable that we found no evidence of admixture even though these populations visually appeared intermediate in plumage between *S. varia* and *S. occidentalis*. Range expansion simulations suggest that we should predict asymmetric introgression into *S. varia* even when the hybridization rate is less than 2% (Currat and Excoffier 2011). Coupled with these predictions, our findings suggest that, although hybridization between *S. varia* and *S. occidentalis* occurs, it has either been vanishingly rare on the edge of the *S. varia* expansion wave or other processes, such as selection or migration, are effectively removing introgressed genetic material from *S. varia* and *S. occidentalis* populations.

We estimated that *Strix varia* has more than ten times greater nucleotide diversity than *S. occidentalis* and we calculated a high *F_ST_* between the species (Table 2), closely matching results from high-coverage genomes of the two species (Hanna *et al.* 2017c). We estimated similar levels of nucleotide diversity in the Siskiyou *S. varia* population and the population comprised of other western *S. varia*, which was consistent with our having found no difference in spotted owl ancestry between these populations (Tables S4 and S5). Similarly, *S. occidentalis* populations pre and post-contact with *S. varia* exhibited similar levels of nucleotide diversity, appeared weakly differentiated, and did not differ in spotted owl ancestry.

We were surprised to find similar levels of nucleotide diversity in western and eastern North American *Strix varia* populations. We expected western *S. varia* populations to harbor lower genetic diversity than the eastern *S. varia* after having been subjected to successive founder effects and corresponding reductions in nucleotide diversity (Austerlitz *et al.* 1997). Simulations have suggested that long-distance dispersal by individuals of a species undergoing a range expansion can inhibit the loss of genetic diversity in the newly formed populations on the edge of the range (Ray and Excoffier 2010). Engler et al. (2016) suggested that this explains why some populations retained genetic diversity in an Old World warbler, *Hippolais polyglotta*, experiencing a range expansion. Recent simulations have also suggested that long-distance dispersal in an invading taxon can counteract introgression of local genetic material into the invader by inhibiting the “surfing” of introgressed genetic regions (Amorim *et al.* 2017). Livezey (2009b) reported the mean natal dispersal distance of *Strix varia* as 41.3 km, but mentioned that some individuals have dispersed as far as 488.1 km. Even if long-distance dispersal has only been occurring at low levels during the *S. varia* range expansion, this could account both the lack of reduction in genetic diversity in western *S. varia* and for the lack of large-scale introgression of *S. occidentalis* genetic material into western *S. varia* populations (Ray and Excoffier 2010; Amorim *et al.* 2017). Long-distance dispersal would have been especially capable of countering introgression of *S. occidentalis* material if non-introgressed *S. varia* were dispersing to the front of the expansion wave (Amorim *et al.* 2017). Long-distance dispersal may also lead to high rates of intraspecific gene flow in western *S. varia*, which could both maintain *S. varia* genetic diversity and counter introgression of *S. occidentalis* genetic material (Ray *et al.* 2003; Currat *et al.* 2008; Petit and Excoffier 2009).

Although our results provide genomic confirmation that hybridization and backcrossing does occur, we found no evidence of widespread admixture between *Strix varia* and *S. occidentalis* in western North America. The distinctive plumage of the *S. varia* individuals collected in Siskiyou County, California, (Figure 1 and Figure S1) does not appear to be a result of hybridization with *S. occidentalis*. We conclude that some plumage characteristics that appear intermediate between *S. varia* and *S. occidentalis* do not in fact indicate hybridization. Previous investigators have issued similar cautionary statements after their genetic studies of hybridization in these taxa (Haig *et al.* 2004; Funk *et al.* 2007). The lack of spotted owl ancestry in these oddly plumaged western *S. varia* suggests that some western *S. varia* may be undergoing drift or local selection, which has affected plumage and size. Coupled with demographic studies (Kelly *et al.* 2003; Dugger *et al.* 2015; Diller *et al.* 2016), our results indicate that the expansion of *S. varia* into the range of *S. occidentalis* in western North America is following a pattern of pure replacement, rather than inducing extinction through hybridization and introgression (Rhymer and Simberloff 1996). It seems unlikely that even introgressed remnants of the *S. occidentalis* genome will remain in areas in contact with *S. varia* if *S. occidentalis* is not able to persist.

## Additional information and declarations

### Author contributions

Conceptualization, Z.R.H., J.P.D., R.C.K.B., J.D.W.; Data Curation, Z.R.H., J.P.D., R.C.K.B.; Formal Analysis, Z.R.H., J.D.W.; Funding Acquisition, Z.R.H., J.P.D., R.C.K.B., J.D.W.; Investigation, Z.R.H., J.D.W.; Methodology, Z.R.H., J.P.D., J.D.W.; Project Administration, Z.R.H.; Resources, Z.R.H., J.P.D., R.C.K.B., J.D.W.; Software, Z.R.H., J.B.H., J.D.W.; Supervision, J.P.D., R.C.K.B., J.D.W.; Validation, Z.R.H.; Visualization, Z.R.H., J.D.W.; Writing – Original Draft Preparation, Z.R.H.; Writing – Review & Editing, Z.R.H., J.P.D., J.B.H., R.C.K.B., J.D.W.

## Acknowledgments

For access to specimens and genetic samples, we thank Lowell Diller; J. Mark Higley and Aaron Pole of the Hoopa Valley Indian Reservation Tribal Forestry department; Susan Haig, Tom Mullins, and Mark Miller of the USGS Forest and Rangeland Ecosystem Science Center; Mary Estes and Morgan of the Chintimini Wildlife Center, Corvallis; Jami Ostby-Marsh and Oroville of the West Valley Outdoor Learning Center, Spokane Valley; Melanie Piazza and WildCare, San Rafael; Sharon Birks and the Burke Museum; Herman L. Mays, Jr., Jane MacKnight, Lauren Hancock, and the Cincinnati Museum Center; Irby Lovette and the Cornell University Museum of Vertebrates; Maureen Flannery, Laura Wilkinson, and the California Academy of Sciences; and Carla Cicero, Theresa Barclay, Shelby Medina, Elizabeth Wommack, and the Museum of Vertebrate Zoology. We thank Anna Sellas for assistance with laboratory work. We generated genomic libraries at the Center for Comparative Genomics, California Academy of Sciences.

## Supporting Information

**Figure S1. Image of barred owls from Siskiyou County.** This image displays the ventral plumage of three *Strix varia* collected in Siskiyou County, California. Owl A is specimen CAS:ORN:92981. Owl B is CAS:ORN:92979. Owl C is CAS:ORN:97181. Author Z.R.H. took this photograph.

**Figure S2. Plot of outlier window proportion versus outlier window length for each sample.** The x-axis plots the lengths of outlier windows in increments of 50,000 nucleotides (nt). The y-axis displays the number of outlier windows of a given length as a proportion of all analyzed windows. The z-axis separates individual samples, which we grouped by population.

**Figure S3. Plot of number of outlier windows versus analyzed windows.** The number of spotted owl ancestry windows of length ≥50,000 nt that were outliers relative to the genome-wide average ancestry for those samples is on the y-axis. The x-axis represents the total number of windows analyzed for each sample. We required the presence of data for at least ten variant sites in order to analyze a window for a given sample. Samples with lower sequence coverage tended to have fewer windows that could be analyzed.

**Table S1. Specimen institution data.** This table provides information regarding the collections that archive the *Strix* specimens utilized in this study.

**Table S2. Additional specimen data.** We here provide additional data for each sample, including the taxonomic identification, the county and state of the collection locality, and the date of collection. The column “Pre or Post Contact” documents whether, based upon the date of collection, a sample’s population was in contact with the other species.

**Table S3. Population assignment, ancestry, and site coverage values for each sample.** The “Category” column provides the population into which we grouped each sample. The spotted owl ancestry values are averages of the ancestry at each variant site across all sites with data for an individual. The site coverage is the average sequence coverage across all sites examined. “SD” stands for “standard deviation”.

**Table S4. Mean and standard deviation spotted owl ancestry by population.** We provide the mean and standard deviation of spotted owl ancestry for each population. The “All Western Barred Owls” population was a superset of the Siskiyou and Western Barred Owl populations. The “All Barred Owls” population is a combination of all of the *Strix varia* samples and the “All Spotted Owls” population is a combination of all of the *S. occidentalis* samples. “SD” stands for “standard deviation”.

**Table S5. Tests of significant difference in spotted owl ancestry.** We here provide the t-values from multiple Welch’s *t*-tests conducted for comparisons of spotted owl ancestry among populations. The “All Western Barred Owls” population was a superset of the Siskiyou and Western Barred Owl populations. An asterisk (*) and bold font indicate those tests with p<0.0125, which is the significance cut-off after applying the Bonferroni correction. The “All Barred Owls” population is a combination of all of the *Strix varia* samples and the “All Spotted Owls” population is a combination of all of the *S. occidentalis* samples.

**File S1. Supplementary details of materials and methods.**

